# A risk stratification approach for improved interpretation of diagnostic accuracy statistics

**DOI:** 10.1101/080366

**Authors:** Hormuzd A. Katki, Mark Schiffman

**Affiliations:** Division of Cancer Epidemiology and Genetics, National Cancer Institute, National Institutes of Health, DHHS, Bethesda, MD, USA

**Author notes:** Corresponding author: **HAK:** Division of Cancer Epidemiology and Genetics, National Cancer Institute, 9609 Medical Center Dr. Room 7E592, Bethesda MD 20892.

## Abstract

Diagnostic accuracy statistics, including predictive values, risk-differences, Youden’s index and Area Under the Curve (AUC), assess the promise of novel biomarkers proposed as diagnostic tests. We reinterpret these statistics in light of risk-stratification (how well a biomarker separates those at higher risk from those at lower risk) to better understand their implications for public-health programs. We introduce an intuitively simple statistic, Mean Risk Stratification (MRS): the average change in risk (pre-test vs. post-test) revealed for tested individuals. High MRS implies better risk separation achieved by testing. MRS demonstrates that conventional predictive values can mislead because they do not account for disease prevalence or test-positivity rates. Little risk-stratification is possible for rare diseases, demonstrating a “high-bar” to justify population-based screening. Importantly, we demonstrate that the risk-difference, Youden’s index, and AUC measure only multiplicative relative gains in risk-stratification: AUC=0.6 achieves only 20% of maximum risk-stratification (AUC=0.9 achieves 80%). However, large relative gains in risk-stratification might not imply large absolute gains if disease is rare or if the test is rarely positive. We illustrate MRS by our experience comparing the performance of cervical cancer screening tests in China vs. the USA. The test with the worst AUC=0.72 in China (visual inspection with ascetic acid) provides twice the risk-stratification of the test with best AUC=0.83 in the USA (human papillomavirus and Pap cotesting) because China has three times more cervical precancer/cancer. MRS could be routinely calculated to better understand the clinical/public-health implications of standard diagnostic accuracy statistics.

## Introduction

After a new biomarker is convincingly associated with disease, the next question is whether it might have clinical/public-health use as a predictive test. Proof of clinical/public-health utility requires absolute (not relative) risks for test results, a valuable risk-reducing intervention, and a comprehensive cost-effectiveness analysis that balances benefits, harms, and costs. Upon biomarker discovery, information required for a comprehensive analysis is usually unavailable. But it remains important to assess preliminarily, without considering costs, benefits, or harms, whether the new biomarker is predictive enough to justify formal cost-effectiveness analyses.

Standard metrics reported for binary biomarkers^1^ provide at best indirect information about predictiveness for clinical/public-health use. The odds-ratio is a well-known poor measure of predictiveness^2^. When comparing two tests, it is uncommon for one test to have both higher sensitivity and specificity, or both higher positive predictive value (PPV) and lower complement of the negative predictive value (cNPV). Two summary statistics, Youden's Index^3^ and the Area Under the Receiver Operating Characteristic Curve (AUC) statistic (AUC is also used for continuous tests)^4,5^, have been correctly criticized for not taking predictive values (i.e. absolute risks) into account, and for not permitting differential weighting of false-positive versus false-negative errors^6^. The AUC is the probability that someone with disease has higher risk than someone without disease, which requires only the risk ranks, not the absolute risks themselves^4^. Because the AUC for a binary test is the average of 1 and Youden's Index^7^, key criticisms of the two tests are shared.

There is a need to better understand the implications of standard diagnostic accuracy metrics for clinical/public-health utility. We reinterpret standard metrics in light of a novel framework for quantifying risk-stratification. Risk-stratification quantifies the ability of a test to separate those at high risk of disease from those at low risk, allowing clinicians to intervene only for those that testing indicates are more likely to develop disease, and not intervene for those that testing indicates will likely not develop disease. We introduce two new broadly applicable, linked statistics that have proven useful in our epidemiologic work on identifying potentially useful biomarker tests for cervical cancer screening. We define mean risk-stratification (MRS) as the average change in risk of disease that a test reveals for tested individuals. We also define a complementary statistic, the Number Needed to Test (NNTest), which quantifies how many people require testing to identify one more disease case than random selection would. We use MRS and NNTest to demonstrate that disease prevalence and test-positivity are crucial for evaluating risk-stratification and interpreting standard metrics. In particular, becaue the maximum MRS is proportional to disease prevalence, little risk-stratification is possible for rare diseases, demonstrating a “high-bar” to justify population-based screening. The novel statistics help place the risk-difference, Youden’s index and AUC into perspective: they measure multiplicative relative gains in risk-stratification, which might not imply large absolute gains if disease is rare. Thus AUC cannot be used to rank tests between populations with different disease prevalence, and we show examples from our experience. Our webtool calculates MRS/NNTest (http://analysistools-dev.nci.nih.gov/biomarkerTools).

## Background for Examples: Cervical cancer screening tests

Human papillomavirus (HPV) causes almost all cervical cancer^8^, leading to development of prophylactic vaccines and HPV DNA testing for screening. HPV DNA-based screening is starting to replace cervical cytology (“Pap smears”), but many new tests are available or being rapidly developed^9–10^. For low/middle-income countries, visual inspection with acetic acid (VIA), a simple low-cost, but unreliable and inaccurate test has been proposed^11,12^. To expedite cervical screening guidelines development^13^, we propose using MRS/NNTest to better interpret standard diagnostic accuracy statistics to identify potentially useful biomarkers for further definitive cost-effectiveness evaluations.

To illustrate the use of MRS/NNTest, we present data from 2 collaborations. Colleagues in China evaluated 3 cervical screening tests (HPV testing, cervical cytology, and VIA) in a previously unscreened population of 30,371 women, to select a test as the basis for a future nationwide screening program.^14^ Second, to support the development of US cervical screening guidelines, we previously analyzed data on 1.4 million women screened since 2003 in the USA at Kaiser Permanente Northern California (KPNC) with cervical cytology, HPV testing, or both concurrently (“cotesting”)^15^.

Table 1 shows the standard metrics for each test in China and KPNC (USA). When comparing two tests in the same population, there is usually a dilemma, namely, a tradeoff of sensitivity vs. specificity, or PPV vs. cNPV, which makes it hard to draw firm conclusions on the basis of a single statistic.

**Table 1.** Characteristics of cervical screening tests in two populations: an unscreened population in China (1.6% precancer/cancer prevalence)^16^ and a previously heavily screened population in the USA at Kaiser Permanente Northern California (0.55% precancer/cancer prevalence)^17^.Acronyms: HPV (human papillomavirus), Pap (Papanicolaou Test), KPNC (Kaiser Permanente Northern California), VIA (visual inspection with acetic acid), PPV (positive predictive value), cNPV (complement of negative predictive value), AUC (Area under the receiver operating characteristic curve), MRS (Mean Risk Stratification), NNTest (Number Needed to Test)

| Population | Cervical screening testing modality | Odds ratio | Test positivity | PPV | cNPV | Sensitivity | Specificity | Youden's Index | AUC | MRS | NNTest |
| --- | --- | --- | --- | --- | --- | --- | --- | --- | --- | --- | --- |
| Unscreened population in China | VIA | 11 | 10.7% | 8.3% | 0.83% | 55% | 90.1% | 45% | 72% | 1.43% | 140 |
|  | HPV | 206 | 17.3% | 9.1% | 0.05% | 98% | 84.0% | 82% | 91% | 2.60% | 77 |
|  | Pap | 108 | 16.7% | 9.4% | 0.10% | 95% | 84.6% | 80% | 90% | 2.60% | 77 |
| Screened population KPNC (USA) | HPV/Pap Cotesting | 37 | 7.5% | 5.4% | 0.16% | 74% | 92.9% | 67% | 83% | 0.73% | 275 |
|  | HPV | 47 | 5.1% | 7.6% | 0.17% | 70% | 95.3% | 65% | 83% | 0.71% | 280 |
|  | Pap | 13 | 3.8% | 4.7% | 0.36% | 34% | 96.3% | 30% | 65% | 0.32% | 633 |

## Methods: Mean Risk-stratification and Number Needed to Test

In the absence of test results or other pre-test information, each individual can only be assigned as a best guess the same population-average risk. After taking a test, people learn how their predicted individual risks differ from population-average. Tests are most useful when they reveal that risk is far enough from population-average risk to justify a change in management.

Denote disease risk in the population as *P(*D*+)* and the fraction testing positive as *P*(*M*+). The positive predictive value is *PPV*=*P(D*+|*M+)*. The complement of negative predictive value is *cNPV*=*P*(*D*+|*M−)*.

Mean Risk-Stratification (MRS) is a weighted average of the increase in risk among those who test positive (i.e., *PPV* – *P(D+)*) and the decrease in risk among those who test negative (i.e., *P(D+)* – *cNPV*):

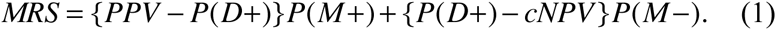

MRS is the average difference between predicted post-test individual risk and population-average risk. Stated simply, MRS is the *average* change in risk that a test reveals for tested individuals. It is easy to show that the two terms on the right-hand side of MRS are equal. Thus

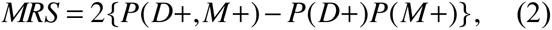

where *P(D+,M+)* is a joint probability. The latter form shows that MRS assesses departure from independence akin to Pearson’s correlation, the Mantel-Haenszel test^16^, and Lewontin’s D’^17^. The **eAppendix** relates MRS to many statistics, including discretized versions of statistics used for continuous biomarkers^18–22^.

An inverse expression of MRS is also useful. A test is only useful if it is substantially better than random selection. Random selection will identify cases by sheer luck: *P(D+,M+)* would equal *P(D+)P(M+)*. The Number Needed to Test (NNTest), to identify 1 more disease case than random selection is

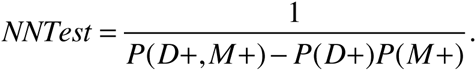

Therefore, 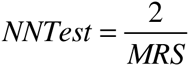. Although MRS and NNTest convey the same information, they are on different scales. MRS is on the scale of absolute risk, allowing comparison to absolute risks of other diseases. NNTest interprets test performance in terms of numbers of people tested, which may help with considering the benefits, harms, and costs associated with the test and subsequent interventions.

For example, HPV testing in an unscreened population in China^14^ changed the risk of cervical precancer/cancer on average by MRS=2.6%, requiring NNTest=77 women to test to identify 1 more precancer/cancer (Table 1). At KPNC, a heavily screened low-risk population in the USA^15^, HPV testing had MRS=0.71%, requiring NNTest=280. MRS conveys how much less *informative* cervical screening, including HPV testing, is in the USA vs. China, due to lower prevalence of disease due to previous screening and treatment. NNTest conveys how much less *efficient* HPV testing is in the US vs. China. More broadly, MRS/NNTest demonstrate the inherent limits of screening for uncommon diseases like cancer: although the few individuals testing positive may have large change in risk, the average change in risk following testing all individuals (the big majority of whom test negative) is modest. Consequently, hundreds may require screening to identify only one more disease case.

The factor of 2 in MRS merits explanation. 1/MRS is the number needed to test to identify, on average, 1 different *outcome* than random selection: 0.5 more disease cases and thus also 0.5 fewer non-disease cases. Thus 2/MRS is the number needed to test to identify 1 more disease case and thus also 1 fewer non-disease case, which is a more natural scaling.

For a perfect test (PPV=1 and cNPV=0), the equation for MRS (1) reveals that the maximum MRS is 50%, which occurs when disease prevalence and test positivity are both 50%. The maximum MRS is not 100% because even if the PPV yields an increase of nearly 1, then the cNPV can yield only a small decrease, and the two are averaged. MRS can be negative (indicating that test-positivity and negativity should be interchanged) and thus MRS is always in the range of [−50%, 50%]. The minimum NNTest is 4 (occuring when disease prevalence and test positivity are both 50%) because random selection tests positive for 2 people out of 4, of which 1 will have disease, thus finding half the 2 disease cases by chance. NNTest can also be negative (≤ −4), indicating that test-positivity and negativity should be interchanged.

When cross-sectional population 2×2 table data are available (not case-control counts), MRS/NNTest can be calculated simply using the cross-product *difference*, or the determinant, of the probabilities in the interior of the 2×2 table. Denoting *a=P(D+,M+), b=P(D+,M−), c=P(D-,M+), d=P(D−,M−)*, and substituting *P(D+)=a+b* and *P(M+)=a+c* into MRS equation (2):

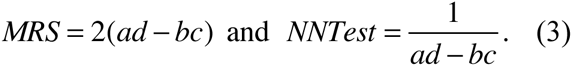

The odds ratio is the cross-product *ratio ad/bc*, hence it is dimensionless. In contrast, MRS is on the scale of risk-differences, and NNTest represents a number of people. In particular, if *a,b,c,d* represent cell counts (rather than probabilities) with *n* being their sum then

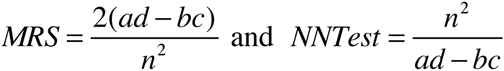

Note that *n*^2^ does not cancel out as it does in the odds ratio. If *n* is the sample size and *a,b,c,d* are probabilities (not counts), the asymptotic variance of MRS can be calculated with the delta-method^16^ on a quadrinomial likelihood for 2×2 tables:

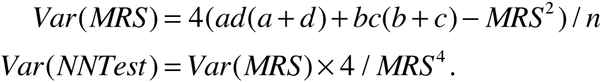

## Results: Implications of MRS and NNTest for evaluating diagnostic tests

### 1. Tests with excellent PPV and cNPV provide little risk-stratification if the tests are rarely positive

Intuitively, risk-stratification seems like it depends only on how far the absolute risks, the PPV and cNPV, spread apart: the risk-difference *t*=*PPV−cNPV*. Substitute into the MRS equation (1) *P*(*D*+) = *PPV* × *P*(*M* +) + *cNPV* × *P*(*M* −), and denoting *p*=P(*M*+), yields:

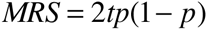

and thus 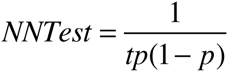. Risk-stratification depends on not only the spread between absolute risks (the risk-difference), but also test-positivity.

A rarely positive test, no matter how big the risk-difference, provides little risk-stratification. For example, consider the hypothetical use of cervical cytology (contrary to usual practice to make a point), dichotomized simply as cancer or not cancer ("not cancer" includes all precancerous and negative cytology results). At KPNC, the 5-year PPV for finding a precancer or cancer, following a cytology result of cancer, is 84%, the cNPV is 0.519%, and the overall precancer/cancer prevalence is 0.524%^23^. The risk-difference *t*=84%-0.519%=83.5% is enormous, suggesting high risk-stratification. However, the probability of having a cytology result of cancer is only 0.006%. Consequently, only with probability 0.006% does a woman get the dramatic risk increase *PPV-P(D+)*=84%-0.524%=83.5%. Almost all the time (99.994%), she has a trivial risk decrease *P(D+)-cNPV*=0.524%-0.519%=0.005%. The MRS is 0.01%: only 1 extra precancer/cancer on average will be found over 5 years per 10,000 women using this test versus random selection. The NNTest=20,086 is an enormous number of women to screen to improve upon random selection. Thus good risk-stratification requires both a sufficiently large risk-difference and a test that is positive sufficiently often.

Figure 1 plots the relationship of MRS/NNTest to the risk-difference for 3 test positivity rates. When risk-difference is 1, the maximum MRS and minimum NNTest are achieved. The importance of test positivity is illustrated by noting that, the MRS/NNTest achieved for risk-difference of 1 is also obtained for a risk-difference of approximately only 0.1 when the test is 10 times as positive (dashed line). Thus a perfect marker for a rarely positive test provides the same risk-stratification as a much weaker-associated marker that is 10 times as positive.

**Figure 1.**
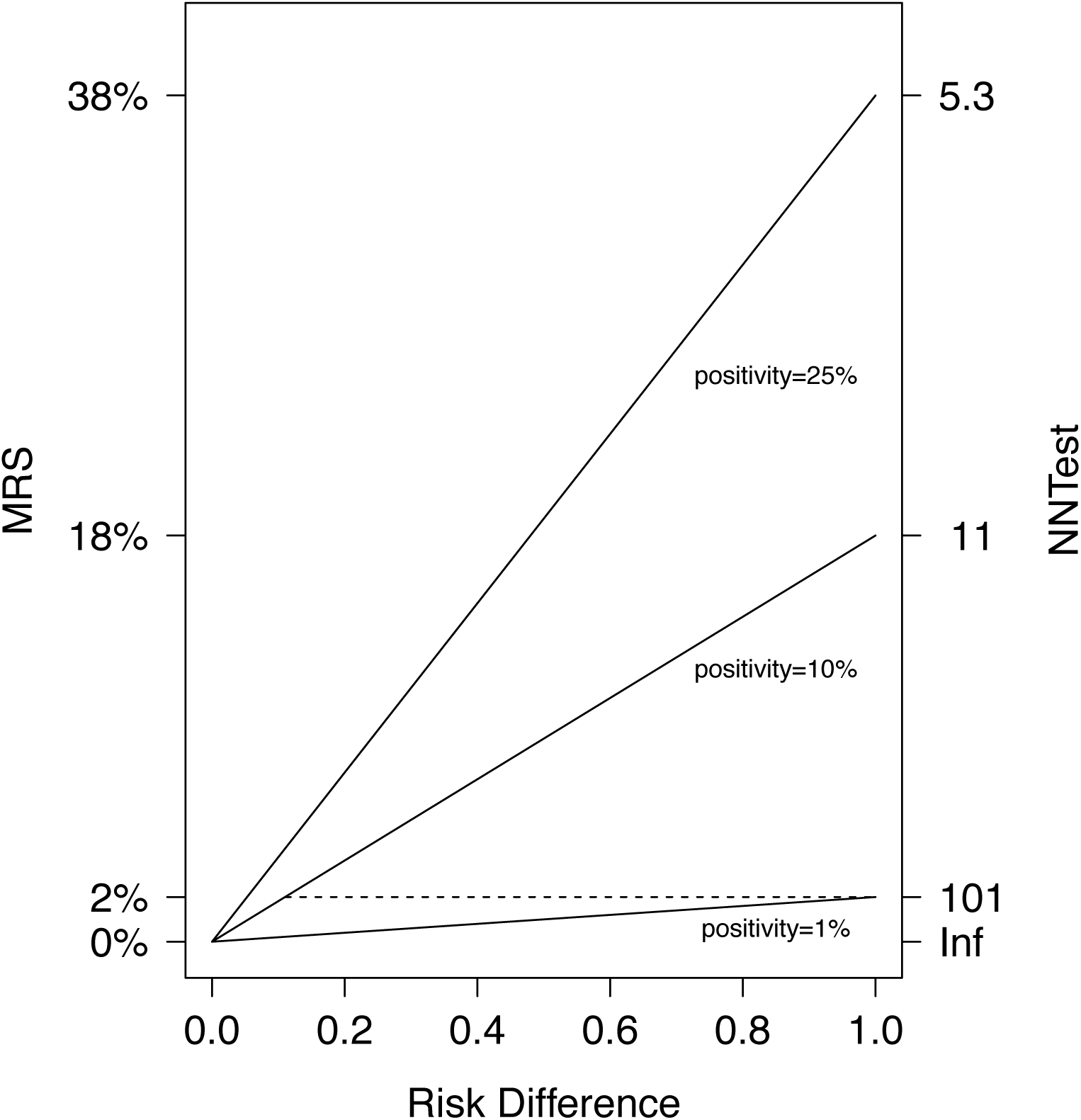
Risk-difference vs. MRS and NNTest for 3 test positivities (1%, 10%, and 25%). Acronyms: MRS (Mean Risk Stratification), NNTest (Number Needed to Test)

### 2. AUC and Youden’s index do not fully measure risk-stratification, because risk-stratification depends strongly on disease prevalence

Neither Youden’s index *J*=*Sens*+*Spec−1* nor AUC depend on absolute risk, but are related to MRS/NNTest. Writing each joint probability in MRS equation (3) as a function of sensitivity and specificity (let *p*\*=*P(D+)*):

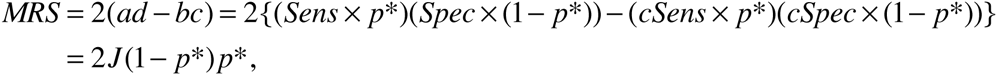

and thus 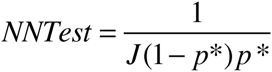. For binary tests, the AUC is the average of sensitivity and specificity^5^, thus substitute *J* = 2 × *AUC* − 1 to get MRS/NNTest in terms of AUC. Thus MRS/NNTest are functions of Youden’s index or AUC and disease prevalence. High AUC or Youden’s index do not imply good risk-stratification if the disease is too rare. Thus:

#### • Disease prevalence defines the maximal amount of risk-stratification that a test can provide

Setting J=1 as a perfect biomarker, the maximum MRS for a disease is *2P(D+)P(D-)*; minimum NNTest is *1/(P(D+)P(D-))*. The rarer the disease, the less risk-stratification is possible. If the maximum risk-stratification is small, every test must inevitably have low risk-stratification This is a reality of general-population cancer screening: on average, screening tests for uncommon diseases such as cancer do not reveal much risk information to tested individuals, and thus, hundreds require screening to identify one more case than would be found by random selection.

In Table 1, the same tests have greater MRS (and lower NNTest) in an unscreened higher-risk population (China; 1.6% disease prevalence) than a screened low-risk population (USA KPNC; 0.55% disease prevalence). The maximum possible MRS is 3 times greater in China vs. KPNC (MRS: 3.2% vs. 1.1%; minimum NNTest: 64 vs. 183, respectively). A test with a fixed Youden’s index (or AUC) should stratify risk better in populations with more disease, such as unscreened populations or populations enriched for high-risk people (such as referral populations), because the maximum possible risk-stratification is greater.

Figure 2 plots the relationship of MRS/NNTest to AUC for 3 uncommon disease prevalences. The importance of disease prevalence is illustrated by noting that, the maximum MRS/NNTest (achieved for AUC=1) is also obtained if AUC=0.55 when disease is 10 times as prevalent (dashed line); AUC=0.6 is required if disease is 5 times as prevalent. Thus a perfect marker for a rare disease provides the same risk-stratification as a weakly associated marker for a disease that is 5 or 10 times as prevalent.

**Figure 2.**
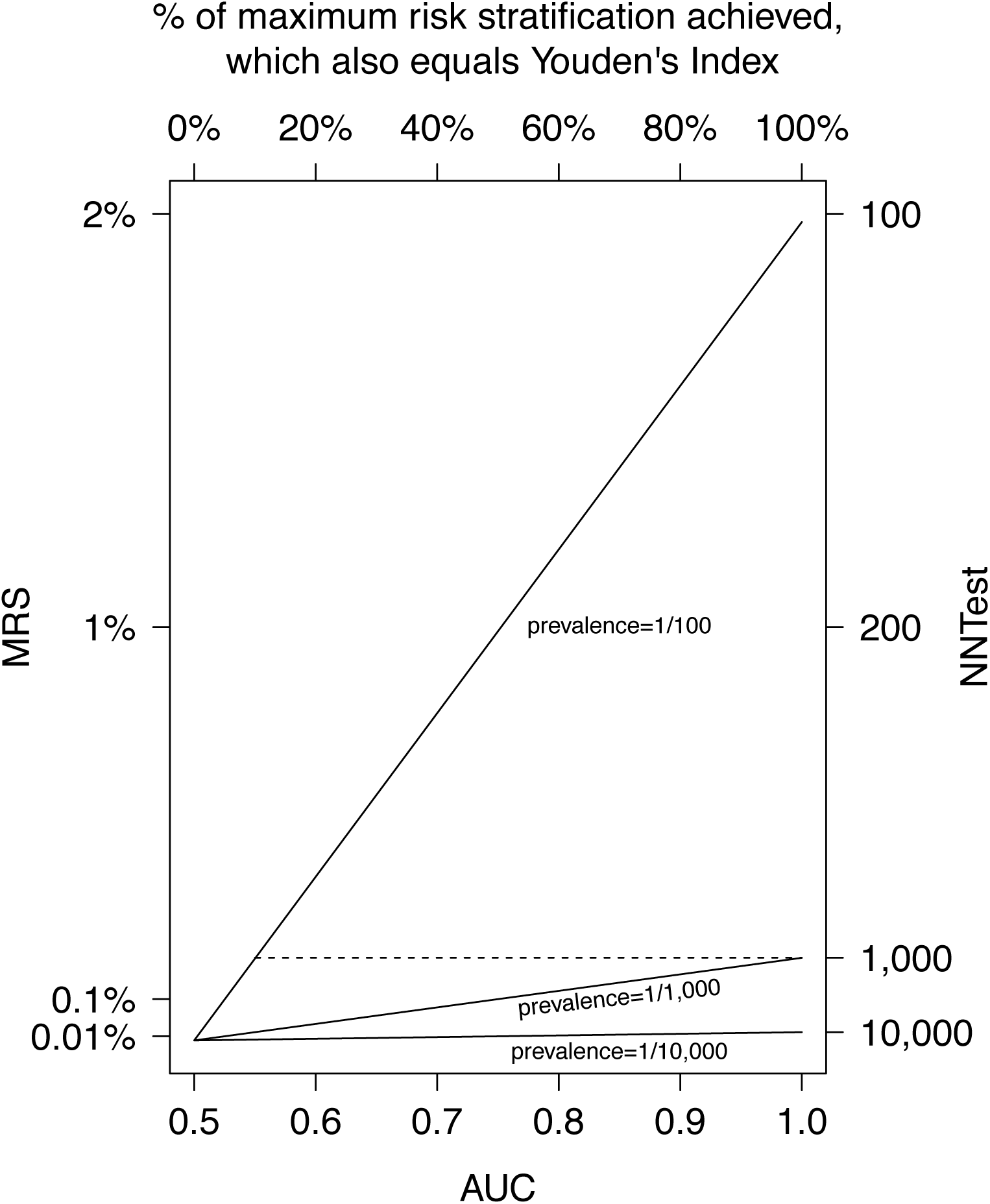
AUC vs. MRS and NNTest for 3 disease prevalences (1/100, 1/1,000, and 1/10,000). Acronyms: AUC (Area under the receiver operating characteristic curve), MRS (Mean Risk Stratification), NNTest (Number Needed to Test)

#### • The risk-stratification implications of Youden's index or AUC depend on disease prevalence

A test with high Youden’s index or AUC in a population with low disease prevalence may provide less risk-stratification than a test with lower Youden’s index or AUC in a population with higher disease prevalence.

In Table 1, cotesting at KPNC in the USA has a greater AUC than VIA in China (83% in KPNC vs. 72% in China), but VIA in China has a better MRS (0.7% vs. 1.4%) and NNTest (275 vs. 140) than cotesting at KPNC. Cotesting is incontestably more accurate than VIA^12^, but VIA stratifies risk better in an easier setting (unscreened population) than cotesting stratifies risk in a harder setting (heavily screened population). Thus, conducting even VIA in China could identify more treatable precancer (and hence prevent more cervical-cancer death) than cotesting at KPNC. Figure 2 shows that for fixed AUC, the MRS/NNTest are dramatically better as disease becomes more prevalent.

#### • Youden's index is the percent of maximum risk-stratification actually achieved by the test. This helps us interpret the meaning of increases in AUC in terms of risk-stratification

Youden’s index (J) is the ratio of the MRS to the maximum MRS achievable for the disease in that population:

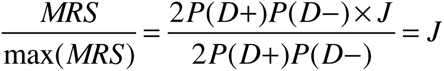

Therefore Youden's index, instead of measuring risk-stratification, measures the percent of maximum risk-stratification actually achieved. This reiterates that Youden’s index and AUC can be high, yet conceal that there is little risk-stratification for uncommon diseases.

This expression also helps interpret the sometimes obscure meaning of increases in AUC^24,25^. Since *J* = 2 × *AUC* − 1, each 1% increase in AUC implies a 2% increase in MRS (or decrease in NNTest). Thus MRS doubles from AUC=0.6 to 0.7. An AUC=0.6 is widely considered to be “modest”, and indeed, only 20% of the maximum MRS is achieved. An AUC=0.7 is widely considered “good”, but only 40% of the maximum MRS is achieved. An AUC=0.9 is required to achieve 80% of the maximum MRS. The top axis of Figure 2 shows how the percent of maximum MRS increases linearly with AUC. As disease becomes rarer, such as for prevalence of 1/10,000, there is little discernible increase in MRS as AUC increases.

Similarly, this expression interprets the risk-stratification implied by an odds-ratio. The minimum odds-ratio required to achieve a Youden’s index has sensitivity equal to specificity. Table 2 shows the minimum odds-ratio required to achieve each fraction of the maximum MRS (the maximum odds-ratio is always infinity, when sensitivity or specificity are 100%). For example, a marker with odds-ratio of 3.4 can attain no higher than 30% of the maximum MRS. If specificity must be high (say 95%), then OR=3.4 can attain no higher than 10% of the maximum MRS. Seemingly large odds-ratios do not suffice to imply large risk-stratification. The odds-ratios in Table 1 follow this pattern. At KPNC, the 3 tests with the lowest and similar MRS of about 0.3% have odds-ratios varying widely (13-36). Similarly, the 4 best tests in China have similar MRS (2.6%-2.8%), but odds-ratios vary widely (108-206). Thus odds-ratios reveal little about risk-stratification.

**Table 2.** The minimum odds-ratios required for a marker to account for each fraction of the maximum possible risk-stratification. The right column provides the minimum odds-ratio if specificity must be high (set at 95%).

| % of maximum risk-stratification achieved | minimum odds-ratio required |  |
| --- | --- | --- |
|  | overall | fix specificity=95% |
| 0% | 1.0 | 1.0 |
| 10% | 1.5 | 3.4 |
| 20% | 2.3 | 6.3 |
| 30% | 3.4 | 10.2 |
| 40% | 5.4 | 15.5 |
| 50% | 9.0 | 23.2 |
| 60% | 16.0 | 35.3 |
| 70% | 32.1 | 57.0 |
| 80% | 81.0 | 107.7 |
| 90% | 361.0 | 361.0 |
| 100% | Infinity | Infinity |

#### • It is important to distinguish rankings of different tests refering to the same population or between populations. Either can be useful depending on the objective

MRS/NNTest and J/AUC will rank the risk-estimation of tests in the same order in the same population because disease prevalence is fixed (see Table 1) – in Figure 1, the MRS increases monotonically with AUC. Thus comparing the MRS/NNTest for 2 tests in the same population is equivalent to comparing their J/AUCs, which is useful for hypothesis testing. However, when prevalence varies across populations, J/AUC do not necessarily even rank test risk-stratification correctly according to MRS/NNTest. For example, a test with AUC=1 has less risk-stratification than a test with AUC>0.55 for a disease 10 times as prevalent (Figure 2, dashed line). Consequently, comparing the risk-stratification of the same test among populations with different prevalence requires MRS/NNTest to account for differing disease prevalence across populations.

#### • MRS and NNTest could be calculated by combining an estimate of Youden’s index (or AUC) in one study with disease prevalence from a target population

MRS/NNTest can be calculated by combining an estimate of Youden’s index from case-control data with disease prevalence from a target population, to quickly assess the risk-stratification implications of a new biomarker in the target population. However, Youden’s index for a new test might not be transportable between populations. For example, Youden’s index and AUC for a given test or test combination are larger in China than at KPNC (Table 1). In China, cervical cytology maintains sensitivity because the women in China are generally unscreened and longstanding prevalent lesions are obvious. In KPNC, cervical cytology has low sensitivity because obvious lesions have been removed by prior screening.

### 3. MRS/NNTest for a rare disease is determined by the difference between specificity and test-negativity

For a rare disease, *MRS* = 2*J*(1 − *p*\*)*p*\* ≈ 2*Jp*\* and 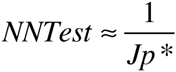. For a rare disease, NNTest is the inverse of Youden’s index scaled by disease prevalence. These approximations are easily rewritten as *MRS* ≈ 2(*Spec* – *P*(*M*–)) and 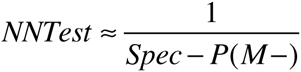. Although specificity is acknowledged as the most important quality of biomarkers for rare diseases^26^, specificity can be artificially boosted by increasing test-negativity. MRS/NNTest balances specificity and test-negativity. Also *Spec* – *P*(*M* –) represents the increase in specificity versus random selection (for random selection, specificity equals test-negativity). In Table 1, the test at KPNC with the best MRS/NNTest is cotesting, which has the lowest specificity but also has the highest sensitivity and the lowest test-negativity at 92.5%. The minimal required test specificity is set by test-negativity. For example, 95% of US women are HPV-negative. Thus HPV tests must have specificity >95% in the US, and 96% specificity is required to achieve an NNTest of 100 in the US. As a rule of thumb, attaining an NNTest of 100 requires that specificity exceed test-negativity by 1%.

## Discussion

After a biomarker-disease association is confirmed, assessment of the predictive capacity of the biomarker is necessary to preliminarily understand the clinical/public-health implications of the biomarker. We introduced two linked metrics, MRS and NNTest, for quantifying risk-stratification from binary diagnostic tests. MRS is the average change in risk that a test reveals for an individual patient. NNTest is the number needed to test to identify 1 more disease case than random selection. We presented 3 major findings on using MRS/NNTest to better understand the clinical/public-health implications of standard diagnostic accuracy statistics.

First, MRS/NNTest demonstrate that there is little risk-stratification possible for rare diseases or for rarely positive tests. Thus screening for rare diseases requires a strong justification. China has 3 times the precancer/cancer prevalence of KPNC in the USA, and all tests, including VIA, the weakest cervical screening test, provide far more risk-stratification in China than at KPNC.

Second, MRS/NNTest demonstrate that the risk-difference, Youden’s index, and AUC measure multiplicative relative gains in risk-stratification, which might not imply large absolute gains if disease is too rare or if the test is too rarely positive. An AUC=0.6 achieves only 20% of maximum risk-stratification. An AUC=0.9 is required to achieve 80%. These findings are generally in accord with intuition that AUC appropriately casts ‘pessimism’ on the power of risk prediction ^24^. Within a population, Youden’s index and AUC will rank tests for risk-stratification in the same order as MRS/NNTest, although rankings may differ across populations with differing disease prevalence.

Third, for uncommon diseases, MRS/NNTest provide a sufficient criterion that ensures high risk-stratification: a high difference between specificity and test-negativity. Thus MRS/NNTest emphasizes the dominant role of specificity, but penalizes for artificially increasing specificity at the expense of test-negativity. Although researchers tend to focus on sensitivity when designing tests, focusing on the difference between specificity and test-negativity would optimize risk-stratification.

This first paper presents the concept of MRS/NNTest. We will extend MRS/NNTest to continuous outcomes to quantify risk-stratification from risk-prediction models and to compare MRS/NNTest to new statistics for evaluating risk models, such as decision curves^27^. Research on estimating MRS/NNTest from different study designs and statistical models is important. For uncommon diseases like cancer, the MRS is small and NNTest is large, and more experience is necessary to develop an intuitive sense of sufficient MRS/NNTest in various settings.

Although etiologic epidemiology progresses using association statistics, standard diagnostic accuracy statistics used in applied epidemiology and public-health are easy to misinterpret. There is a torrent of new biomarkers and risk-prediction models, but evaluating them remains challenging, and many are used clinically without sufficient formal study^28^. The STARD guidelines^1^ require reporting sensitivity, specificity, predictive values, or AUC, but none of these directly informs about risk-stratification and the odds-ratio reveals little about risk-stratification. Thus MRS/NNTest might be worth adding to studies of test performance to quantify risk-stratification and better understand the meaning of standard statistics. For example, it would be immediately clear from MRS/NNTest why *BRCA1/2* testing of all women, justified on the basis of high PPV and risk-difference^29^, would have little public-health value because of low mutation prevalence (0.25%). However, MRS/NNTest would also immediately clarify that restricting testing to populations with higher mutation prevalence, such as Ashkenazi Jews (2.5%), is more likely to have public-heath value. MRS/NNTest can help temper premature claims of clinical/public-health utility but also help justify future definitive studies that account for all necessary components (benefits, harms, and costs) to decide clinical/public-health usefulness. Our risk-stratification webtool is publicly available (http://analysistools-dev.nci.nih.gov/biomarkerTools).

## Acknowledgements

We acknowledge our late friend, mentor, and collaborator, Dr. Sholom Wacholder, who made seminal contributions to this work. We thank Holly Janes for her comments on a earlier version of this paper. We thank our colleagues at Kaiser Permanente Northern California and in China for their collaboration.

